# White Matter Extension of the Melbourne Children’s Regional Infant Brain Atlas: M-CRIB-WM

**DOI:** 10.1101/590596

**Authors:** Sarah Hui Wen Yao, Joseph Yuan-Mou Yang, Bonnie Alexander, Michelle Hao Wu, Claire E. Kelly, Gareth Ball, Marc L. Seal, Peter J. Anderson, Lex W. Doyle, Jeanie L.Y. Cheong, Alicia J. Spittle, Deanne K. Thompson

**Author notes:** **Corresponding Author Information:** Joseph Yuan-Mou Yang, Murdoch Children’s Research Institute, Royal Children’s Hospital, Flemington Road, Parkville, Victoria (3052), Australia.

## Abstract

**Background:** Understanding typically developing infant brain structure is crucial in investigating neurological disorders of early childhood. Brain atlases providing standardised identification of neonatal brain regions are key in such investigations. Our previously developed Melbourne Children’s Regional Infant Brain (M-CRIB) and M-CRIB 2.0 neonatal brain atlases provide standardised parcellation of 100 and 94 brain regions respectively, including cortical, subcortical, and cerebellar regions. The aim of this study was to extend the M-CRIB atlas coverage to include 54 white matter regions.

**Methods:** Participants were ten healthy term-born neonates who comprised the sample for the M-CRIB and M-CRIB 2.0 atlases. WM regions were manually segmented on *T*_2_ images and co-registered diffusion tensor imaging-based, direction-encoded colour maps. Our labelled regions are based on those in the *JHU-neonate-SS* atlas, but differ in the following ways: 1) we included five corpus callosum subdivisions instead of a left / right division; 2) we included a left / right division for the middle cerebellar peduncle; and 3) we excluded the three brainstem divisions. All segmentations were reviewed and approved by a paediatric radiologist and a neurosurgery research fellow for anatomical accuracy.

**Results:** The resulting neonatal WM atlas comprises 54 WM regions: 24 paired regions, and six unpaired regions comprising five corpus callosum subdivisions and one pontine crossing tract. Detailed protocols for manual WM parcellations are provided, and the M-CRIB-WM atlas is presented together with the existing M-CRIB and M-CRIB 2.0 cortical, subcortical and cerebellar parcellations in ten individual neonatal MRI datasets.

**Conclusion:** The updated M-CRIB atlas including the WM parcellations will be made publicly available. The atlas will be a valuable tool that will help facilitate neuroimaging research into neonatal WM development in both healthy and diseased states.

## 1. Introduction

Parcellated brain atlases are a key component of many neuroimaging tools. They can facilitate identification and labelling of brain regions in a consistent manner, such that properties of these regions can be compared across brains, and across time points. Until recently, few parcellated atlases were available for the crucial neonatal time period where the foundations for all future neurodevelopment are set. During the neonatal period, MRI images have relatively low spatial resolution due to small brain size, and have different tissue contrast compared with older children and adults due to partial myelination and dynamic tissue properties in neonates (Heemskerk et al., 2013). Over the last decade, increasing efforts in the neonatal brain imaging field have led to development of several neonatal parcellated atlases (Alexander et al., 2019; Alexander et al., 2017; Blesa et al., 2016; de Macedo Rodrigues et al., 2015; Gousias et al., 2012; Kuklisova-Murgasova et al., 2011; Makropoulos et al., 2016; Oishi et al., 2011; Shi et al., 2010; Shi et al., 2011). These atlases differ in image modality and quality, parcellation technique, and parcellation schemes. Many of these atlases were defined on *T*_2_-weighted images (which provide higher tissue contrast than *T*_1_-weighted images due to partial myelination at the neonatal time point) and focus on parcellation of cortical regions and deep grey nuclei. White matter (WM) segmentation has generally been provided as a single label or a few regions (Alexander et al., 2019; Alexander et al., 2017; de Macedo Rodrigues et al., 2015), or included together with adjacent grey matter (GM) in parcellated regions (Gousias et al., 2012; Shi et al., 2011; Tzourio-Mazoyer et al., 2002). The major white matter tracts are extant at term (Dubois et al., 2014), however these cannot be defined based on *T*_*1*_-or *T*_*2*_-weighted images alone. In order to delineate anatomical tracts within WM, diffusion weighted images (DWI), which provide information about WM fibre orientation, are required.

One atlas to date, the ‘*JHU-neonate-SS*’ atlas (Oishi et al., 2011) has provided manually delineated anatomical WM regions using neonatal DWI data. The atlas consists of voxel-wise averaging of 122 parcellated brain regions altogether, including 52 WM regions, from the MRI data of 25 healthy neonates. The WM regions were manually segmented based on a single participant’s MRI dataset, which was then warped to the group-averaged brain template. The parcellated detail of the *JHU-neonate-SS* atlas is unprecedented, and demonstrates the ability of the included regions to be delineated at term. Importantly, the *JHU-neonate-SS* atlas provides standardised identification of regions at the neonatal time point. However, using atlas labels based on a single individual does not allow individual variability in brain morphology to be captured (Alexander et al., 2017; Mori et al., 2008; Wang et al., 2014).

Accounting for individual anatomical variance is an important factor to consider when studying a period of brain development that is marked by significant brain structural changes and growth (Shi et al., 2011). Some endeavours have been made to address this issue by warping existing single-subject atlases to multiple neonatal subjects, with the aim of providing larger atlas training sets with greater inter-subject variability. For example, Shi et al. (Shi et al., 2011) warped the adult parcellated ‘*Automated Anatomical Labelling*’ (AAL) atlas (Tzourio-Mazoyer et al., 2002) (defined based on *T*_1_ images) to an infant longitudinal sample. However, it is generally acknowledged that warping adult atlases to infant space offers limited accuracy, due to morphological differences between the adult brain and the developing neonatal brain (Alexander et al., 2017; Blesa et al., 2016; Dickie et al., 2017; Fillmore, Richards, Phillips-Meek, Cryer, & Stevens, 2015; Kazemi, Moghaddam, Grebe, Gondry-Jouet, & Wallois, 2007; Richards, Sanchez, Phillips-Meek, & Xie, 2016; Sanchez, Richards, & Almli, 2012). Additionally, warping a single parcellated image to multiple participants, even those of the same age, is likely to introduce labelling error related to imperfect registration aligning different target brains. This occurs in instances where there are individual differences in morphology or image properties between the template and the target brains (Akhondi-Asl, Hoyte, Lockhart, & Warfield, 2014). Target brains that differ more greatly from the template will incur more marked registration error, and thus a training set that captures some of the individual variability in the population is valuable. The ‘gold standard’ procedure for defining an accurate and broadly applicable parcellated atlas is manual segmentation in a large sample of representative individuals (Gousias et al., 2012; Shi et al., 2010).

We previously presented the *Melbourne Children’s Regional Infant Brain* (M-CRIB) (Alexander et al., 2017) and M-CRIB 2.0 (Alexander et al., 2019) multi-subject (*N* = 10), manually parcellated whole-brain atlases. The M-CRIB and M-CRIB 2.0 parcellations include cortical regions that are compatible with the *Desikan-Killiany* (Desikan et al., 2006) and *Desikan-Killiany-Tourville* (Klein et al., 2012) adult cortical parcellations, respectively. Compatibility of neonatal atlases with those commonly used at older timepoints is important for longitudinal investigations of brain development and tracking the progression of developmental disorders (de Macedo Rodrigues et al., 2015; Gousias et al., 2012; Oishi et al., 2011). There is currently an unmet need for a multi-subject, manually parcellated neonatal WM atlas to provide standardised identification of WM regions in a way that is compatible with atlases commonly used at older time points. The aim of this study was to extend the coverage of the M-CRIB atlases to include manually parcellated WM regions by utilising neonatal DWI data. We elected to model our parcellation scheme on that provided by the *JHU-neonatal-SS* atlas (Oishi et al., 2011) which has label nomenclature and terminology consistent with the commonly used adult JHU atlas (Mori et al., 2008; Mori et al., 2005; Oishi et al., 2009) In this paper, we detail a manual WM parcellation scheme in ten term-born neonates, presented as a WM extension of the M-CRIB atlases, which we have named the M-CRIB-WM atlas.

## 2. Materials and Methods

### 2.1 Participants

Participants were ten healthy term-born neonates (≥37 weeks’ gestation; four females; gestational age at scanning 40.29 - 43.00 weeks, *M* = 41.71, *SD* = 1.31). These participants were the same sample utilised for our existing M-CRIB atlases (Alexander et al., 2017; Alexander et al., 2019). The participants were initially selected from a larger cohort of control infants with MRI scans, recruited as part of preterm studies (Spittle et al., 2014; Walsh, Doyle, Anderson, Lee, & Cheong, 2014) on the basis of minimal motion and other artefact on *T*_2_-weighted images (Alexander et al., 2017). Neonates who received resuscitation at birth, were admitted to a neonatal intensive care or special care unit, had a birth weight of less than 2.5 kg, or had congenital conditions affecting growth and development, were excluded (Spittle et al., 2014; Walsh et al., 2014). All ten participants selected were assessed at age two years and did not have any major health problems, cerebral palsy or major cognitive delay (Alexander et al., 2017).

This study was approved by the Royal Children’s Hospital Human Research Ethics Committees. Informed parental/guardian consent was obtained prior the study commencement.

### 2.2 MRI data acquisition and pre-processing

MRI scans were acquired at the Royal Children’s Hospital, Melbourne, Australia, on a 3-Tesla Siemens MAGNETOM Trio Tim scanner. The neonates were scanned during non-sedated natural sleep. They were first fed, swaddled and fitted with ear plugs and ear muffs throughout the MRI study. Transverse *T*_*2*_ restore turbo spin echo sequences were acquired with 1 mm axial slices, flip angle = 120°, repetition time (TR) = 8910 ms, echo time (TE) = 152 ms, field of view (FOV) = 192 × 192 mm, matrix = 384 × 384, and in-plane resolution 1 mm^2^ (zero-filled interpolated to 0.5 × 0.5 × 1 mm). DWI sequences were acquired using a multi-*b*-value, single-shot echo planar imaging (EPI) sequence with TR = 20400 ms, TE = 120 ms, FOV = 173 × 173 mm, matrix = 144 × 144, 100 axial slices, 1.2 mm isotropic voxels, 45 non-collinear gradient directions, *b*-values ranging from 100-1200 s/mm^2^, and 3 *b* = 0 s/mm^2^ volumes. The total diffusion sequence was divided into three separate acquisitions to improve compliance, and if any of the diffusion acquisitions had unacceptable levels of motion artefact, the scan was repeated whenever possible until acceptable diffusion images were acquired. All infants were scanned with the same diffusion sequence, including the same range of *b*-values.

The *T*_*2*_ images were bias corrected using N4ITK (Tustison et al., 2010), and skull-stripped using the Functional MRI of the Brain (FMRIB) Software Library (FSL) Brain Extraction Tool (BET) (Smith, 2002). They were then aligned to anterior commissure-posterior commissure (AC-PC) line using 3D Slicer, and resampled to 0.63 mm isotropic (preserving voxel volume) using the FMRIB’s Linear Image Registration Tool (FLIRT) (Greve & Fischl, 2009; Jenkinson, Bannister, Brady, & Smith, 2002; Jenkinson & Smith, 2001).

The DWI data were corrected for head motion and eddy current-induced distortions using the FSL ‘eddy_correct’ tool (Jenkinson & Smith, 2001), incorporating *b*-vector reorientation (Leemans & Jones, 2009). Echo planar image distortions were corrected based on a gradient echo field map and FMRIB’s Utility for Geometrically Unwarping Echo planar images (FUGUE), as previously described (Thompson et al., 2018). The diffusion tensor imaging (DTI) model was fitted using the weighted linear least squares method in FSL. The principal DTI eigenvector, representing the modeled single-fibre orientation per voxel, was used to generate a fibre direction-encoded colour (DEC) map with the default colour scheme: the Anterior-Posterior (AP) oriented fibres encoded in green; Left-Right (LR)-oriented fibres encoded in red; and Superior-Inferior (SI) oriented fibres encoded in blue.

### 2.3 Manual WM parcellation methods

All WM parcellation was performed on the *T*_*2*_-weighted and co-registered DWI images in volume space using Insight Toolkit (ITK)-SNAP v 3.6.0 (Yushkevich et al., 2006), which simultaneously displays axial, sagittal and coronal views along with a composite 3D surface representation of utilised labels.

To aid parcellation of regions containing boundaries between multiple adjacent WM tracts, we constructed individual vector maps (Supplementary Data 1) where each voxel was assigned a category based on the angular distance between the principal direction of diffusion and each image axis (AP, LR or IS). These maps were used to clarify the region boundaries in situations where the primary fibre direction was not obvious via visual inspection of the DEC map. This creates artificial boundaries to help distinguish between voxels containing tracts running predominantly e.g. SI adjacent to voxels containing tracts running AP. The *b* = 0 s/mm^2^ images were then nonlinearly registered to the *T*_*2*_ images using Advanced Normalization Tools (ANTs) (Avants, Epstein, Grossman, & Gee, 2008; Avants et al., 2011), and this registration was applied to all the DTI images to bring them into *T*_2_ space.

We overlaid the following parameter maps on *T*_*2*_-weighted structural images in a stepwise manner, in order to remove boundary ambiguity between different WM regions. First, we overlaid the co-registered DEC map, which revealed the principal fibre direction in each WM region. Next, the co-registered vector map was overlaid to aid boundary definition, as voxels were only selected if they had the same vector profile as the principal fibre direction for the WM region. The parcellation boundaries were then checked against the underlying *T*_2_-weighted images to ensure that they conformed with the structural landmarks specified as boundaries.

Parcellation was performed and checked on a combination of axial, sagittal and coronal slices, leveraging the clearest perspective available for each WM structure. All structures were parcellated collectively, brain-by-brain, instead of individually, region-by-region. The visibility of adjacent structures provided much insight in delineating the boundaries of the current parcellated region. Manual parcellation was completed by one operator (S.Y.). A neurosurgery research fellow (J.Y.), and a paediatric radiologist (M.W.) confirmed the accuracy of each region’s boundaries on all brains, based on the proposed parcellation scheme used in this study.

All parcellations were then masked using the M-CRIB WM labels (Fig. 1).

**Figure 1.**
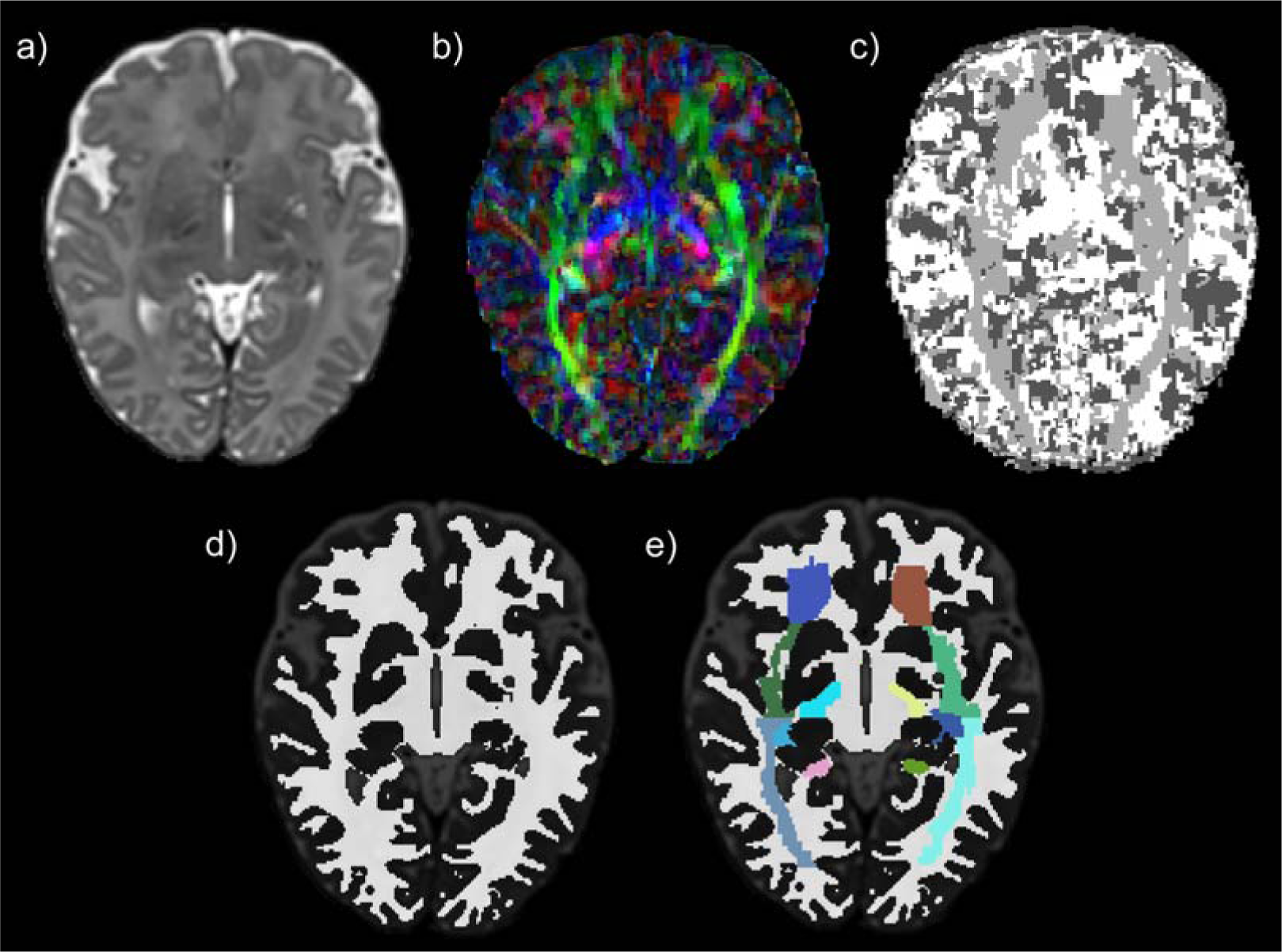
Illustration of the stepwise white matter (WM) parcellation process. (a) *T*_2_– weighted image; (b) Direction encoded colour (DEC) map; (c) Vector map; (d) WM mask; (e) WM mask with parcellated regions overlaid. Images are displayed in radiological orientation.

We derived our parcellation scheme from the JHU neonatal atlas parcellation scheme (Oishi et al., 2011) with modifications to improve anatomical clarity of the following WM structures. First, we incorporated five corpus callosum (CC) subdivisions based on Hofer’s Classification (Hofer & Frahm, 2006), instead of a three-part (genu, body and splenium) and Left-Right (LR) division. Next, we utilised a LR division for the middle cerebellar peduncle (MCP), instead of a singular region for both MCPs. Third, we labelled the pontine crossing tract (PCT) as a singular region instead of a LR division. Lastly, we excluded brainstem divisions in our parcellation scheme.

### 2.4 Protocols for delineating the WM regional boundaries

#### 2.4.1 Regions in the Brainstem

##### Corticospinal Tracts (CST)

*<u>Description</u>:* This parcellation defines the portion of CST in the ventral pons (Ture, Yasargil, Friedman, & Al-Mefty, 2000). They consist of SI-oriented fibres (blue-coloured on DEC map). *<u>Relevant boundaries</u>: <u>Superior:</u>* The midbrain-pontine junction *<u>Inferior</u>:* The pontine-medullary junction *<u>Posterior</u>:* The PCT (described below).

##### Pontine Crossing Tract (PCT)

*<u>Description</u>:* This parcellation consists of LR-oriented fibres from the pontine nuclei in the ventral pons (red-colour on DEC map). *<u>Relevant boundarie</u>s: <u>Superior</u>*: The midbrain-pontine junction. *<u>Inferior</u>:* The pontine-medullary junction. *<u>Anterior</u>:* The CST (described above). *<u>Posterior</u>:* The Medial Lemniscus (ML; described below) (Mori et al., 2008).

##### Medial Lemniscus (ML)

*<u>Description</u>:* This parcellation defines the portion of ML in the ventral pons. They consist of SI-oriented fibres (blue-coloured on DEC map) (Mori et al., 2008). *<u>Relevant boundaries</u>: <u>Superior</u>:* The midbrain-pontine junction. *Inferior:* The pontine-medullary junction. *<u>Anterior</u>:* The PCT (described previous).

##### Superior Cerebellar Peduncle (SCP)

*<u>Description</u>:* This structure contains mainly efferent cerebellar fibres, connecting the cerebellum to the midbrain. It is most easily distinguished from between the level of the cerebellar nuclei and the midbrain using the DEC map (Mori et al., 2008). *<u>Relevant boundaries</u>: <u>Superior</u>:* It is marked by the SCP decussation fibres (red-coloured on the colour DEC map) at the level of the midbrain. (Mori et al., 2008) *<u>Inferior</u>:* The dentate nuclei of cerebellum, located medio-posteriorly to the MCP (described below).

##### Middle Cerebellar Peduncle (MCP)

*<u>Description</u>:* This structure contains entirely afferent cerebellar fibres, connecting the cerebellum to the ventral pons. *<u>Relevant boundarie</u>s: <u>Posterior</u>:* The dentate nuclei in the cerebellum. *<u>Medial:</u>* The CST, PCT and ML parcellations in the ventral pons, best visualised on the axial plane.

##### Inferior Cerebellar Peduncle (ICP)

*<u>Description</u>:* This structure contains the spinocerebellar fibre tracts, connecting the cerebellum to the medulla and spinal cord. *<u>Relevant boundaries</u>: <u>Superior</u>:* The level of the mid-pons. *<u>Inferior</u>:* The dorso-lateral aspect of the medulla (Hirsch et al., 1989).

#### 2.4.2 Projection Fibres

The projection fibres enter or exit the brain via the spinal cord by traversing through the following structures – from inferiorly to superiorly (for ascending afferent fibres; or in reverse order for descending efferent fibres): cerebral peduncle (CP), internal capsule (IC) and corona radiata (CR). The CP-IC boundary was arbitrarily defined at the level of the anterior commissure (Mori et al., 2008). The IC-CR boundary was arbitrarily defined at the axial level where the internal and external capsules merged (Mori et al., 2008). The IC can be identified on axial planes as the “bend-shaped” WM region located between the caudate nucleus, the lentiform nucleus and the thalamus. It was arbitrarily divided into four parts: the anterior limb (ALIC), the genu, the posterior limb (PLIC) and the retrolenticular part (RLIC). The CR was arbitrarily divided into three parts: anterior (ACR), superior (SCR), and posterior corona radiata (PCR) (Mori et al., 2008).

##### Cerebral Peduncle (CP)

*<u>Description</u>:* The IC converges into the CP, forming the ventral portion of the midbrain. *<u>Relevant boundaries</u>: <u>Superior</u>:* The CP-IC plane at the anterior commissure level (Mori et al., 2008). *<u>Inferior</u>:* The midbrain-pontine junction.

##### Anterior Limb of Internal Capsule (ALIC)

*<u>Description</u>:* The ALIC is the anterior bend of the IC in front of the genu. We included the genu of IC into this parcellation – as per the approach adopted by the *JHU-neonatal-SS* atlas. *<u>Relevant boundaries</u>: <u>Superior</u>:* The IC-CR plane against the ACR (Mori et al., 2008). *<u>Inferior</u>:* The CP-IC plane (Mori et al., 2008). *<u>Medial</u>:* The head of the caudate nucleus. *<u>Lateral</u>:* The lentiform nucleus. *<u>Posterior</u>:* The PLIC (described below).

##### Posterior Limb of Internal Capsule (PLIC)

*<u>Description</u>:* This structure represents the posterior bend of the IC, behind the genu. *<u>Relevant boundaries</u>: <u>Superior</u>:* The CR-IC plane against the SCR. *<u>Inferior</u>:* The CP-IC plane. *<u>Anterior</u>:* The genu of the IC. *<u>Posterior</u>:* The RLIC. This posterior boundary can be identified by a change in the dominant fibre orientation identified on the DEC map - from predominantly blue-coloured SI-oriented fibres in the PLIC to predominantly green-coloured AP-oriented fibres in the RLIC. *<u>Medial</u>:* The thalamus. *<u>Lateral</u>:* The lentiform nucleus.

##### Retrolenticular Part of Internal Capsule (RLIC)

*<u>Description</u>:* This portion of the IC is caudal to the lenticular nucleus and carries the optic radiation. *<u>Relevant boundaries</u>: <u>Superior</u>:* The CR-IC plane against the PCR. *<u>Inferior</u>:* The sagittal stratum (SS) at the CP-IC plane. *Anterior:* The PLIC. *<u>Posterior:</u>* The posterior thalamic radiation (PTR). This posterior boundary was arbitrarily defined by an imaginary vertical line extending from the midpoint of the CC splenium in the mid-sagittal plane (Mori et al., 2008).

##### Anterior Corona Radiata (ACR)

*<u>Description:</u>* The CR refers to the subcortical WM region inferior to the centrum semiovale and superior to the IC. *<u>Relevant boundaries: Inferior:</u>* The CR-IC plane against the ALIC (Mori et al., 2008). *<u>Posterior:</u>* The superior corona radiata (SCR). This posterior boundary was defined arbitrarily by an imaginary vertical line extending from the posterior edge of the genu of CC in the mid-sagittal plane (Mori & van Zijl, 2007)

##### Superior Corona Radiata (SCR)

*<u>Description:</u>* Arbitrary boundaries were used to parcellate the SCR. *<u>Relevant boundaries: Inferior:</u>* The CR-IC plane against the PLIC (described previously). *<u>Anterior:</u>* The ACR (described previously). *<u>Posterior:</u>* The PCR. This posterior boundary was defined arbitrarily by an imaginary vertical line extending from the anterior edge of the CC splenium in the mid-sagittal plane (Mori et al., 2008; Mori & van Zijl, 2007)

##### Posterior Corona Radiata (PCR)

*<u>Description:</u>* Arbitrary boundaries were used to parcellate the PCR. *<u>Relevant boundaries: Inferior:</u>* By the RLIC (anteriorly) and the PTR (posteriorly). This was defined arbitrarily by an imaginary horizontal line extending from the midpoint of the CC splenium in the mid-sagittal plane (Mori et al., 2008). *<u>Anterior:</u>* The SCR (described previously). *<u>Medial:</u>* The forceps major and tapetum (TAP; described below) of the CC. *<u>Lateral:</u>* The superior longitudinal fasciculus (SLF; described below).

#### 2.4.3 Association Fibres

##### Cingulum Cingular Part (CGC)

*<u>Description:</u>* This parcellation defines the frontal component of the cingulum WM within the cingulate gyrus. The cingulate gyrus is located immediately above the CC and below the cingular sulcus. It curves around the back of the CC splenium and continues as the hippocampal part of the cingulum (CGH) that enters the mesial temporal lobe (Shah, Jhawar, & Goel, 2012). *<u>Relevant boundaries: Superior</u>*: The cingular sulcus. *<u>Inferior:</u>* The CC. *<u>Posterior</u>:* This was arbitrarily delineated against the CGH by an imaginary horizontal line extending from the midpoint of the CC splenium in the mid-sagittal plane (Mori et al., 2008). This division corresponds to a change in the dominant fibre orientation identified on the DEC map - from predominantly green-coloured AP-oriented fibres in the CGC to predominantly blue-coloured SI-oriented fibres in the CGH. *<u>Lateral:</u>* The CC body fibres.

##### Cingulum Hippocampal Part (CGH)

*<u>Description:</u>* The CGH courses within the parahippocampal gyrus and terminates anteriorly in the mesial temporal lobe (Shah et al., 2012). *<u>Relevant boundaries: Superior:</u>* The CGC (described previously). *<u>Temporal terminations:</u>* We arbitrarily defined this to be at the level of the hippocampal head in the sagittal plane.

##### Fornix (Fx)

*<u>Description:</u>* This parcellation defines the forniceal body and column to the level of the anterior commissure (Nieuwenhuys, 2008; Shah et al., 2012). The pre-commissural column fibres to the septal region were not included due to limited image resolution. The forniceal body can be identified immediately inferiorly to the CC in the mid-sagittal plane. *<u>Relevant boundaries: Superior</u>*: The body of CC. *<u>Anterior and inferior</u>*: At the level of the anterior commissure. *<u>Posterior</u>*: Before the two forniceal bodies diverge to become the forniceal crus.

##### Stria Terminalis (ST)

*<u>Description:</u>* This WM tract is the main efferent fibre pathway from the amygdala that courses along the ventricular surface of the thalamus (Nieuwenhuys, 2008; Shah et al., 2012). This parcellation also includes the forniceal crus and fimbria – differentiation of these two fibre tracts is not possible due to the current limited image resolution. The forniceal crus courses immediately posterior to the thalamus, and medial to the SS (Mori et al., 2008). It continues caudally as the fimbria when entering the mesial temporal lobe (Nieuwenhuys, 2008; Shah et al., 2012). *<u>Relevant boundaries:</u> Anterior and superior*: the forniceal body. *<u>Temporal terminations:</u>* We defined the fimbria terminations arbitrarily at two axial slices above the level of the CP, at the diencephalon-mesencephalon junction.

##### Superior Longitudinal Fasciculus (SLF)

*<u>Description:</u>* This WM tract provides connections to the frontal, parietal and temporal lobes (Martino et al., 2013). It is located dorso-laterally to the CR (Mori et al., 2008). *<u>Relevant boundaries: Medial:</u>* The fronto-parietal component of the SLF (contains predominantly green-coloured AP-oriented fibres) is bounded medially by the CR (contains predominantly blue-coloured SI-oriented fibres). The temporal component of the SLF (contains predominantly blue-coloured SI-oriented fibres) is bounded medially by the PTR anteriorly and the SS posteriorly (both of which contain predominantly green-coloured AP-oriented fibres).

##### External Capsule (EC)

*<u>Description:</u>* This parcellation includes both the external capsule (EC) and extreme capsule, with the intervening GM, the claustrum. Separating these regions is not possible due to limitations in the image resolution. It excludes the portion of the EC containing the inferior fronto-occipital fasciculus (IFO) and the uncinate fasciculus (UFC), both of which are parcellated separately (described below). *<u>Relevant boundaries: Superior:</u>* The axial slice where the EC and IC merge. *<u>Inferior:</u>* An arbitrary boundary against the IFO - identified by the changes in the dominant fibre orientation from the DEC map (from predominantly SI-oriented, blue-coloured fibres in the EC to predominantly AP-oriented, green-coloured fibres in the IFO). *<u>Medial:</u>* The lentiform nucleus and the IC (Mori et al., 2008). *<u>Lateral:</u>* The insular cortex.

##### Posterior Thalamic Radiation (PTR)

*<u>Description:</u>* This parcellation contains WM tracts that connect the caudal thalamus to the occipital and parietal lobes. The fibres are predominantly AP-oriented, green-coloured fibres on the DEC map, and are best visualised on the axial plane. *<u>Relevant boundaries: Superior:</u>* The PCR (described previously). *<u>Inferior:</u>* The SS (described below) (Mori et al., 2008). *<u>Anterior:</u>* The RLIC (described previously).

##### Sagittal Stratum (SS)

*<u>Description:</u>* This WM region contains long association WM tracts, such as the IFO, optic radiation and PTR, with fibre projections to the occipital lobe. Division against the PTR parcellation is arbitrary. The SS was best visualised on the sagittal plane (Mori et al., 2008). *<u>Relevant boundaries: Superior:</u>* The RLIC (anteriorly) and the PTR (posteriorly). This superior boundary was arbitrarily defined at the anterior commissure level. *Anterior:* The IFO (described below) and the UFC (described below).

##### Superior Fronto-Occipital Fasciculus (SFO)

*<u>Description:</u>* This WM tract, typically vestigial in humans, connects the occipital and frontal lobes and extends posteriorly along the dorsal edge of the caudate nucleus (Forkel et al., 2014; Jellison et al., 2004). *<u>Relevant boundaries</u>: <u>Superior and lateral</u>:* The SCR (described previously) (Mori et al., 2008). *<u>Medial</u>:* The head of caudate nucleus and the lateral ventricle.

##### Inferior Fronto-Occipital Fasciculus (IFO)

*<u>Description:</u>* This WM pathway forms the long-ranged frontal and occipital connection (Jellison et al., 2004; Martino, Brogna, Robles, Vergani, & Duffau, 2010; Mori et al., 2008). This parcellation defines the portion of the IFO that traverses through the EC. *<u>Relevant boundaries: Superior:</u>* The EC. *<u>Inferior:</u>* The insular segment of the UFC at the temporal stem (Choi, Han, Yee, & Lee, 2010). Its boundary against the UFC was identified by the changes in the dominant fibre orientation from the DEC map (from predominantly AP-oriented, green-coloured fibres for the IFO to blue-coloured SI-oriented fibres for the UFC) (Ebeling & von Cramon, 1992; Kier, Staib, Davis, & Bronen, 2004) *<u>Posterior:</u>* The SS (described previously) (Jellison et al., 2004). *<u>Medial:</u>* The lentiform nucleus. *<u>Lateral:</u>* The insular cortex.

##### Uncinate Fasciculus (UFC)

*<u>Description:</u>* This hook-shaped WM tract connects the orbitofrontal lobe to the anterior temporal lobe via the temporal stem portion of the EC (Ebeling & von Cramon, 1992; Jellison et al., 2004; Kier et al., 2004). This parcellation defines the insular segment of the UFC that traverses through the temporal stem (Choi et al., 2010). *<u>Relevant boundaries: Superior:</u>* The IFO (described previously). *<u>Inferior:</u>* The midbrain-pontine junction. *<u>Posterior:</u>* The SS (described previously).

#### 2.4.4 Commissural Fibres

The CC provides the main inter-hemispheric connections for the cerebrum. We did not parcellate the anterior and posterior commissures due to their small sizes in neonates and the limited image resolution. We adopted the Hofer’s classification, segmenting the CC into five divisions, based on its AP length at the mid-sagittal plane (Hofer & Frahm, 2006). The TAP contains the CC temporal fibres and was parcellated as a separate region. The CC is best located in the mid-sagittal plane, above the lateral ventricles and below the cingulate gyrus. The CR marks the lateral boundaries for the CC parcellations (Mori et al., 2008).

##### Corpus Callosum I (CCI)

*<u>Description:</u>* The CCI is the anterior 1/6 of the CC at the mid-sagittal plane. It represents both the rostrum and the genu of the CC (Hofer & Frahm, 2006).

##### Corpus Callosum II (CCII)

*<u>Description:</u>* The CCII is defined as the portion of the CC between the anterior 1/2 and the anterior 1/6 of the CC at the mid-sagittal plane. It represents the anterior body of the CC (Hofer & Frahm, 2006).

##### Corpus Callosum III (CCIII)

*<u>Description:</u>* The CCIII is between the posterior 1/2 and the posterior 1/3 of the CC at the mid-sagittal plane. It represents the posterior body of the CC (Hofer & Frahm, 2006).

##### Corpus Callosum IV (CCIV)

*<u>Description:</u>* The CCIV is between the posterior 1/3 and the posterior 1/5 of the CC at the mid-sagittal plane. It represents the isthmus of the CC (Hofer & Frahm, 2006).

##### Corpus Callosum V (CCV)

*<u>Description:</u>* The CCV is the posterior 1/5 of the CC at the mid-sagittal plane. It represents the splenium of the CC (Hofer & Frahm, 2006).

##### Tapetum (TAP)

*<u>Description:</u>* The TAP is best located within the lateral ventricular wall at the level of ventricular trigone in the occipital lobe. It is medial to the SS, and can be readily distinguished from the SS on the DEC map due to differences in the predominant fibre orientation (TAP contains predominantly blue-coloured, SI-oriented temporal CC fibres; the SS contains predominantly green-coloured, AP-oriented association fibres) (Mori et al., 2008).

### 2.5 Data availability statement

The individual parcellated and structural images of the M-CRIB-WM neonatal atlas, including complete whole-brain atlases comprising M-CRIB and M-CRIB 2.0 regions (basal ganglia, thalamus, cerebellum, cortex, and other regions) and the current WM regions, will be publicly available via https://github.com/DevelopmentalImagingMCRI. Public provision of these datasets has been approved via Murdoch Children’s Research Institute, and is compliant with ethics agreements via the Royal Children’s Hospital Human Research Ethics Committee.

## 3. Results

The WM extension of the M-CRIB atlases comprises 24 pairs of left-and right-hemispheric structures, and six single structures, totalling 54 regions. A full list of parcellated WM regions is included in Table 1. Combining the M-CRIB-WM regions with the original M-CRIB (Alexander et al., 2017) whole-brain atlas, results in a parcellated atlas comprising 154 regions altogether. Combining the M-CRIB-WM regions with the M-CRIB 2.0 (Alexander et al., 2019) whole brain atlas results in 148 parcellated regions. Selected axial slices of the WM parcellations and combined M-CRIB 2.0 and WM parcellations for a single participant are illustrated in Figure 2. Figure 3 depicts a surface representation of all WM parcellations for a single participant. Figure 4 illustrates the surface representation of all WM parcellations for a single neonatal participant compared with that for an adult *T*_*1*_-weighted brain image labelled using the equivalent JHU parcellation scheme (Mori et al., 2008).

**Table 1.**
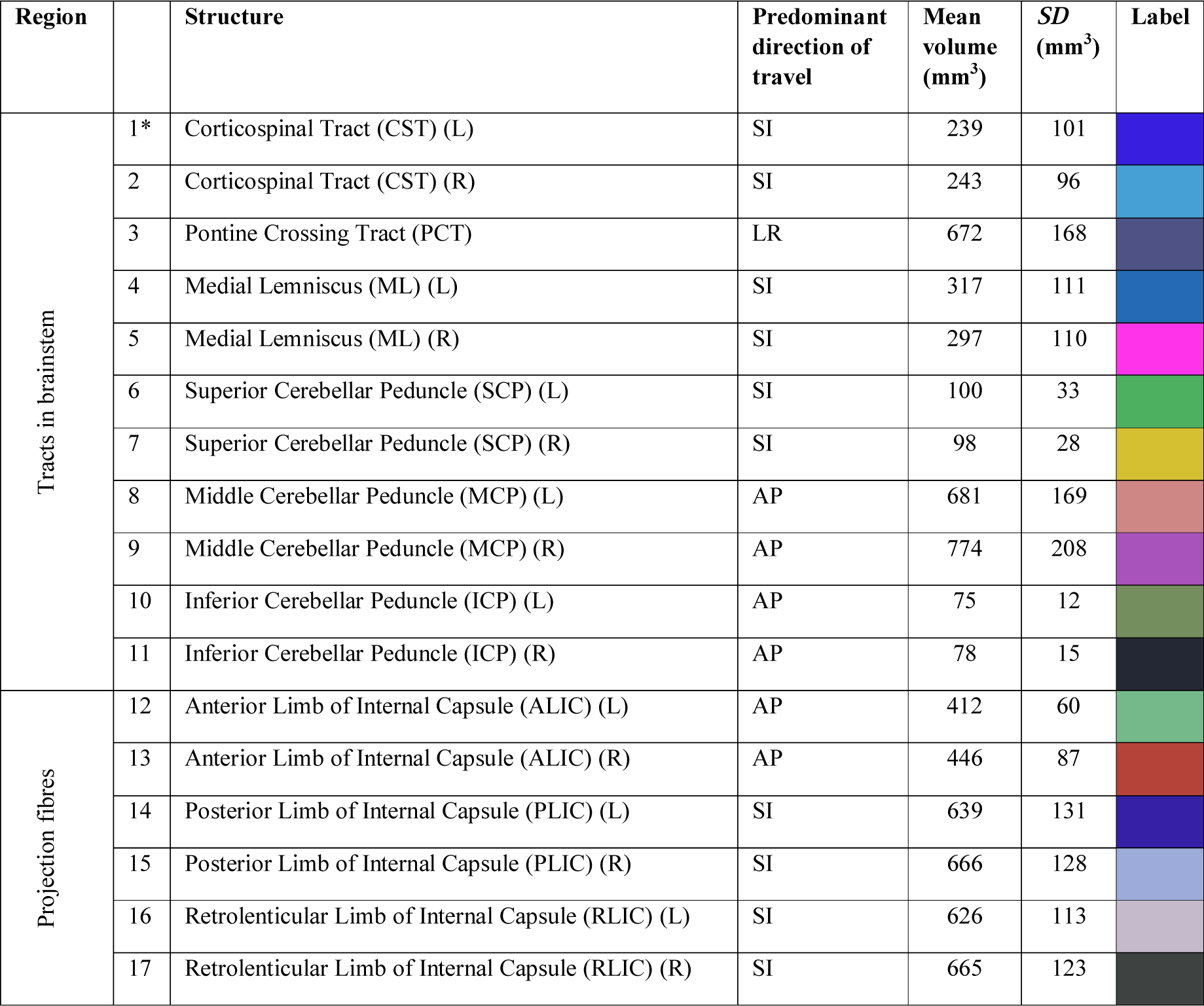
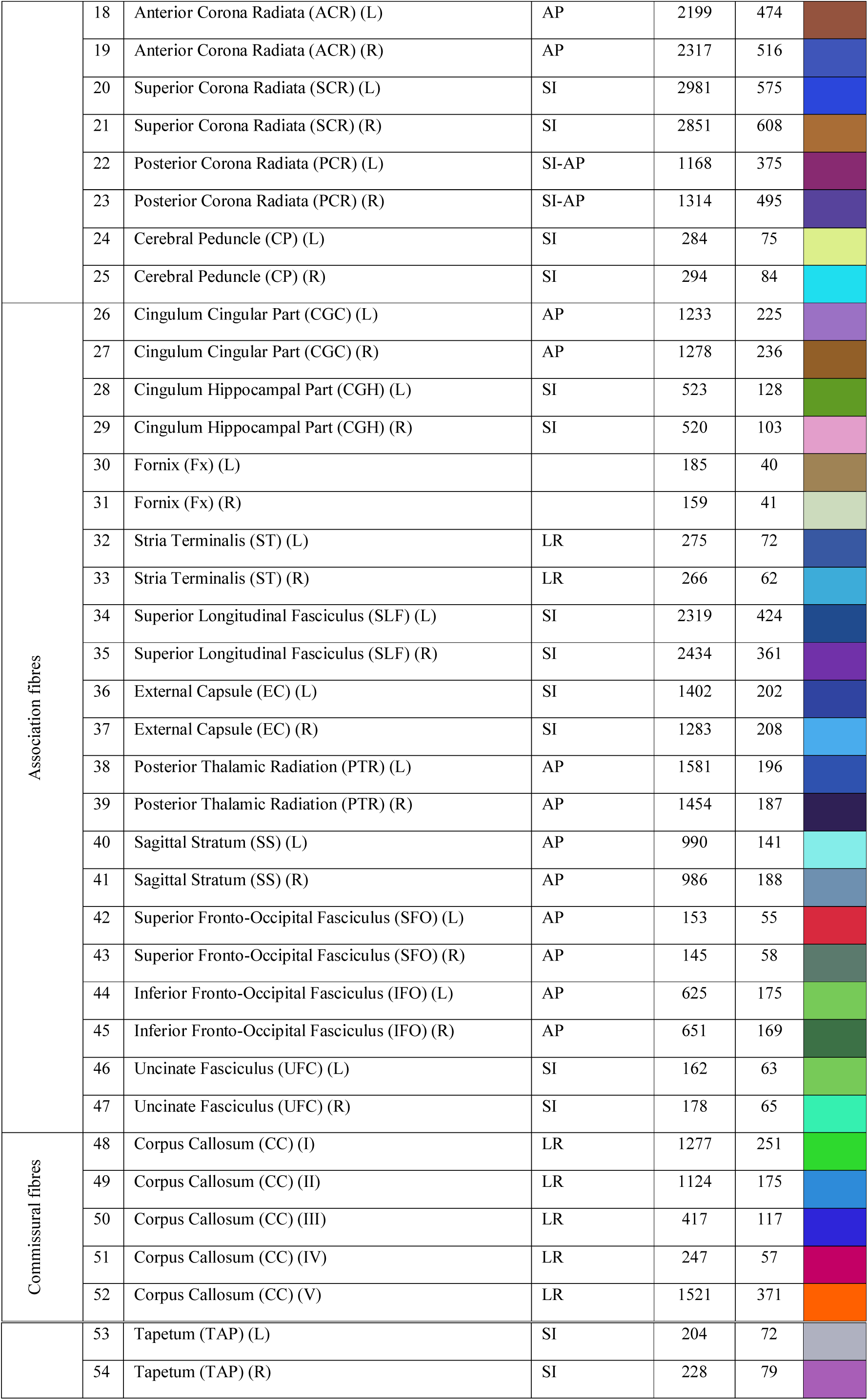
Complete list of parcellated M-CRIB-WM regions and mean volume for each region Note: SI: Superior-Inferior; LR: Left-Right; AP: Anterior-Posterior. *Numbers listed are not label indices. Labels files containing label indices are provided via our GitHub page: https://github.com/DevelopmentalImagingMCRI

**Figure 2.**
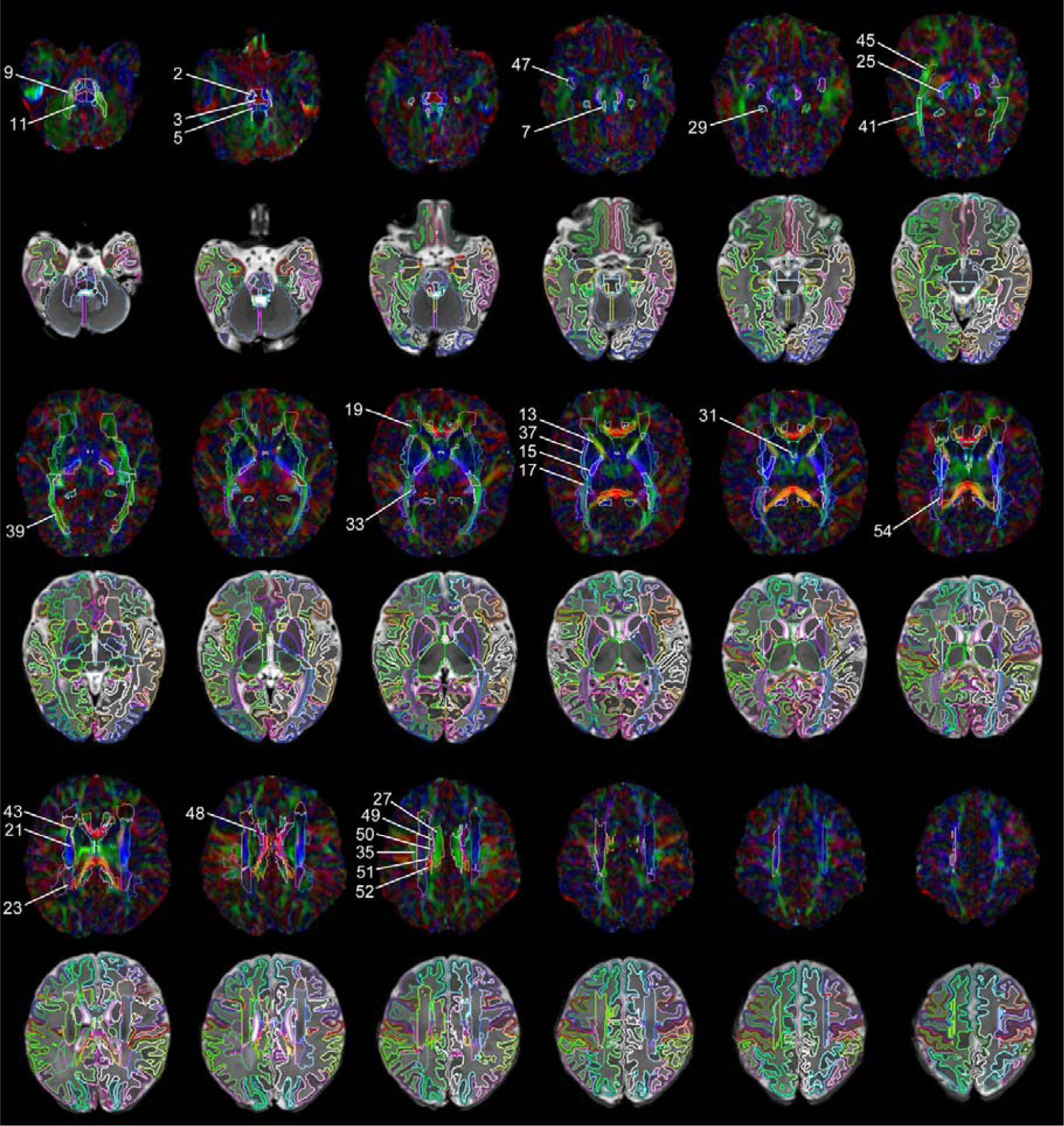
Selected axial slices of M-CRIB-WM white matter (WM) parcellations (1^st^, 3^rd^ and 5^th^ rows) overlaid on DWI images; and M-CRIB-WM parcellations combined with M-CRIB 2.0 atlas GM regions (2^nd^, 4^th^, and 6^th^ rows) overlaid on *T*_2_-weighted images. Images are for a single participant, shown in 5-slice increments. Images are displayed in radiological orientation. Annotated region numbers correspond to those listed in Table 1. For full detail, see the online, high-resolution version of this image.

**Figure 3.**
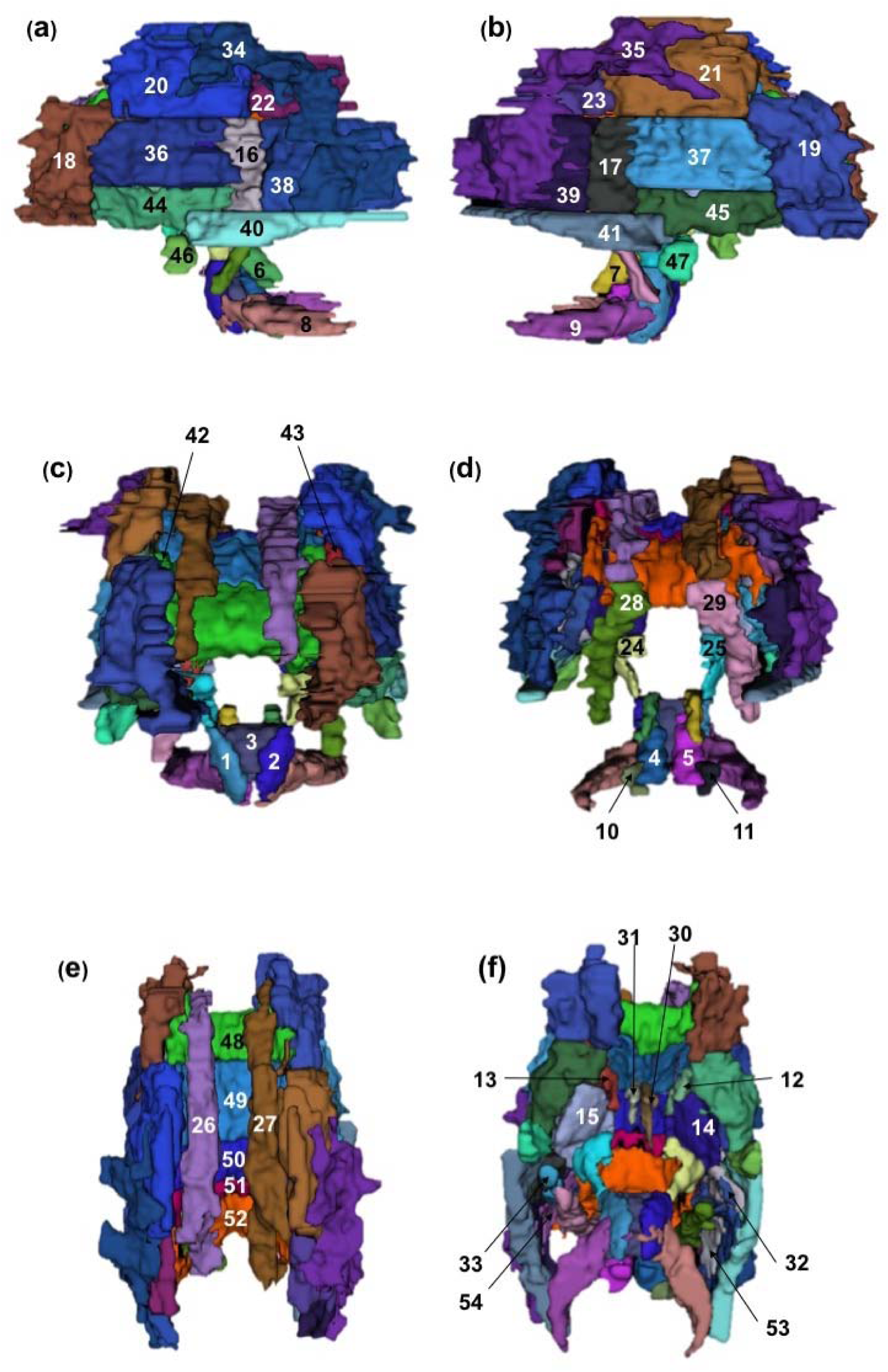
Annotated 3D representation of all white matter (WM) parcellations for a single participant. (a) Left hemisphere; (b) Right hemisphere; (c) Frontal view; (d) Occipital view; (e) Sagittal view; (f) Inferior view. Surfaces underwent Gaussian smoothing with SD 0.8 mm for display purposes. Labels correspond to structures listed in Table 1.

**Figure 4.**
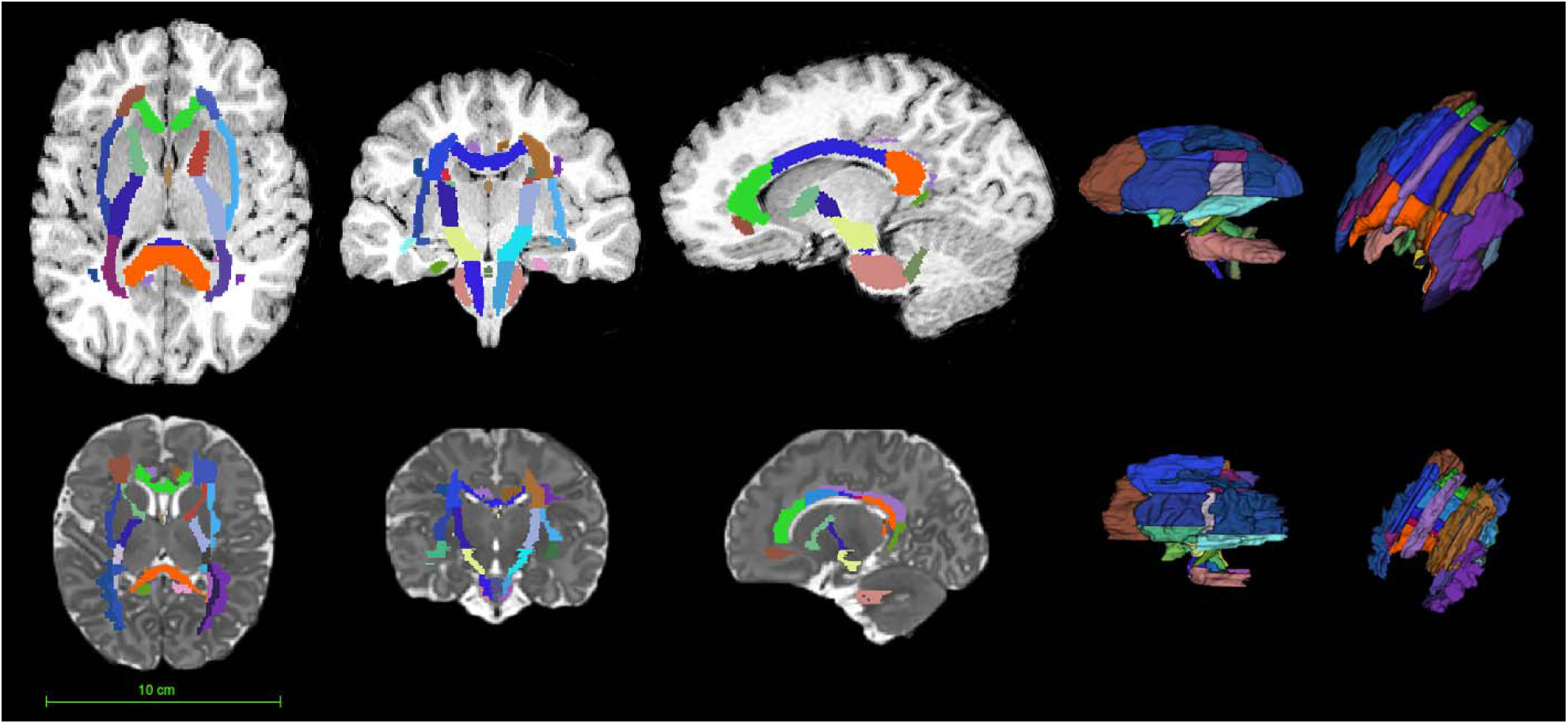
Comparison of the adult JHU white matter (WM) atlas and the M-CRIB-WM atlas. Top row: *T*_1_-weighed image of a healthy 18-year-old brain that has been labelled with the adult JHU atlas (Mori & van Zijl, 2007). Bottom row: *T*_2_-weighed image of a single neonatal participant from the M-CRIB-WM sample, with manually parcellated labels overlaid. Relative size of the adult and neonate brains is to scale.

Supplementary Figure S1 presents surface representations of all ten parcellated WM atlases. Supplementary Figure S2 presents surface representations of combined WM and cortical regions in a single participant.

## 4. Discussion

In this work, we present the WM extension to our existing M-CRIB atlases, the M-CRIB-WM. This atlas contains 54 manually parcellated WM regions, in ten healthy term neonates. The M-CRIB-WM has been defined based on high-quality neonatal DWI and *T*_2_-weighted data, enabling delineation of the relatively small, detailed structures in the neonatal brain. The use of manual segmentation allowed us to precisely segment structures in each individual. This resulted in detailed ground truth parcellations that capture individual variability in morphology, thus allowing this atlas to be more accurately applicable to a larger proportion of the population.

Manual segmentation remains the best practice for MRI brain parcellation as it allows precise delineation of different brain regions, particularly those with complex or arbitrarily defined boundaries. The WM of the brain, comprising a complex network of neuronal axons, typically has indistinct boundaries between neighbouring WM tracts. For example, the AP-oriented fibres of the IFO, optic radiation, and PTR traverse through the SS, and cannot be differentiated macroscopically even with meticulous cadaveric fibre dissection techniques (Yasargil, Ture, & Yasargil, 2004). In other instances, the only discernible feature between neighbouring WM tracts is the difference in the dominant fibre orientation. For example, the IFO and UFC are differentiable in the temporal stem because one (IFO) has AP-oriented fibres, forming the fronto-occipital connections, whereas the other (UFC) has SI-oriented fibres and hooks around the temporal stem, forming the fronto-temporal connections (Choi et al., 2010; Ebeling & von Cramon, 1992; Kier et al., 2004). The WM regions are defined for convenience in anatomical studies. The distinction between neighbouring WM regions is also largely arbitrary. While some WM regions can be defined structurally based on anatomical landmarks (for example, the CP), there are other WM regions where arbitrary boundaries are unavoidable. In particular, the deep WM regions (ACR, SCR and PCR) lack clear, recognisable anatomical boundaries between the neighbouring regions. Here, we used imaginary lines drawn perpendicularly from the back edge of the genu of the CC, and the front edge of the splenium of the CC to divide the three portions of the CR, a technique we developed through visual inspection of the JHU adult brain atlas. Although arbitrary, we found this boundary definition reproducible through all subjects.

Terminology that is anatomically clear can reduce ambiguity and bias, and allow consistency across different operators. We based our region definitions on those provided for the *JHU-neonate-SS atlas*, and have further elaborated boundary definitions for the purpose of clarity. For example, the CST label consists of only the ventral pontine portion of this WM tract in the JHU atlases, rather than the entire tract. The resulting detailed parcellation protocols that we have provided for all 54 WM regions, as an elaboration of those provided for the JHU atlases, are a strength of the M-CRIB-WM atlas.

A challenge when performing manual parcellation based on DWI is that the dominant colour intensities on the DEC map may be ambiguous, particularly in regions where multiple WM tracts intersect. Our approach to this problem was to develop a discretised vector map that indexes only the principle direction (i.e., AP, SI, or LR) in each voxel. This facilitated the identification of a predominant colour on the corresponding DEC map, and thus clarified the principle fibre direction at the boundary between neighbouring regions. This provided a decision solution that enabled us to define anatomical regions with an increased level of certainty.

Considering there are equivalent JHU atlases available for older time points (Mori et al., 2008; Oishi et al., 2009), our compatible neonatal WM atlas also has the benefit of facilitating longitudinal analyses of neuroimaging metrics from equivalent WM regions. Together with the large range of complementary multi-parametric neuroimaging tools and techniques available, the combined M-CRIB and M-CRIB-WM atlases will enable detailed structural and microstructural measures to be obtained in an accurate and age-specific way for major cortical and subcortical regions of the neonatal brain, and now additionally all major WM tracts and regions. To our knowledge, there has not previously existed a single neonatal atlas encompassing the parcellation of both extensive GM and WM regions to a satisfactory level of detail. Our novel M-CRIB-WM atlas, along with the M-CRIB cortical and subcortical atlases, provide neonatal whole brain MRI coverage of standardised GM and WM parcellations. This addition will greatly benefit the field of infant neuroimaging research.

In summary, we have presented a neonatal WM atlas capturing important WM structural variability unique to, and characteristic of, the neonatal time point. The individual parcellated and structural images of the M-CRIB-WM neonatal atlas, and versions combined with the M-CRIB whole-brain atlases, will be publicly available via https://github.com/DevelopmentalImagingMCRI. This novel atlas will provide extensive neonatal brain coverage with substantial anatomic detail. It will be a valuable resource that will help facilitate investigation of brain structure at the neonatal time point, and developmentally across the lifespan.

## List of Abbreviations

AAL: Automated Anatomical Labelling
ACR: Anterior Corona Radiata
ALIC: Anterior Limb of Internal Capsule
ANTs: Advanced Normalization Tools
AP: Anterior-Posterior
BET: Brain Extraction Tool
CC: Corpus Callosum
CGC: Cingulum Cingular Part
CGH: Cingulum Hippocampal Part
CP: Cerebellar Peduncle
CR: Corona Radiata
CST: Corticospinal Tracts
DEC: Direction-Encoded Colour
DTI: Diffusion Tensor Imaging
DWI: Diffusion Weighted Images
EC: External Capsule
EPI: Echo Planar Imaging
FLIRT: Functional Magnetic Resonance Imaging of the Brain’s Linear Image Registration Tool
FMRI: Functional Magnetic Resonance Imaging
FOV: Field of View
FSL: Functional Magnetic Resonance Imaging Software Library
FUGUE: FMRIB’s Utility for Geometrically Unwarping Echo planar images
Fx: Fornix
GM: Grey Matter
IC: Internal Capsule
ICP: Inferior Cerebellar Peduncle
IFO: Inferior Fronto-Occipital Fasciculus
ITK: Insight Toolkit
JHU: Johns Hopkins University
LR: Left-Right
MCP: Middle Cerebellar Peduncle
M-CRIB: Melbourne Children’s Regional Infant Brain
M-CRIB-WM: Melbourne Children’s Regional Infant Brain-White Matter
ML: Medial Lemniscus
MRI: Magnetic Resonance Imaging
PCR: Posterior Corona Radiata
PCT: Pontine Crossing Tract
PLIC: Posterior Limb of Internal Capsule
PTR: Posterior Thalamic Radiation
RLIC: Retrolenticular Part of Internal Capsule
SCP: Superior Cerebellar Peduncle
SCR: Superior Corona Radiata
SFO: Superior Fronto-Occipital Fasciculus
SI: Superior-Inferior
SLF: Superior Longitudinal Fasciculus
SS: Sagittal Stratum
ST: Stria Terminalis
TAP: Tapetum
TE: Echo Time
TR: Repetition Time
UFC: Uncinate Fasciculus
WM: White Matter

## Acknowledgements

We gratefully acknowledge ideas and support provided by the members of the Victorian Infant Brain Study (VIBeS) and Developmental Imaging groups, as well as the Melbourne Children’s MRI Centre, located at the Murdoch Children’s Research Institute, Melbourne, Victoria. We also thank the families who participated in this study. This project was supported by the Australian National Health and Medical Research Council (NHMRC) (Project Grant ID 1028822 and 1024516; Centre of Clinical Research Excellence Grant ID 546519; Centre of Research Excellence Grant ID 1060733; Senior Research Fellowship ID 1081288 to P.J.A.; Early Career Fellowship ID 1053787 to J.L.Y.C., ID 1053767 to A.J.S., ID 1012236 to D.K.T.; Career Development Fellowship ID 1108714 to A.J.S., ID 1085754 to D.K.T.), the Royal Children’s Hospital Foundation (RCH 1000 to J.Y.M.Y), Murdoch Children’s Research Institute, The University of Melbourne Department of Paediatrics, and the Victorian Government’s Operational Infrastructure Support Program. The funding sources had no involvement in the study design; in the collection, analysis and interpretation of data; in the writing of the report; and in the decision to submit the article for publication.

Declarations of interest: none.

## Contributorship Statement

L.D., J.C., A.S., and P.A designed the original cohort studies and acquired the MRI scans. D.T. and J.Y. conceptualized and designed the study. B.A., C.K., and G.B. contributed to the preparation of MRI images utilized in the parcellation process. S.Y. performed the manual parcellations. S.Y., J.Y. and M.W. contributed to development of the parcellation protocols. B.A., J.Y. and S.Y. contributed to the writing of the manuscript. B.A. and S.Y. contributed to the production of figures in the manuscript. All authors revised the manuscript and approved the final version to be submitted.

## Supplementary Material

**Supplementary Data 1.** Script for co-registered vector map used to aid in boundary delineation.

~~~
#! /bin/bash

# diffusion_directions.sh

# creates a 3-class image with classes defined by closest axis to the principal direction of diffusion in each voxel

# usage:
#./diffusion_directions.sh <V1 image> <diffusion_brain_mask>
# eg:./diffusion_directions.sh img_V1.nii.gz img_mask.nii.gz

# output: xyz.nii.gz
# where all voxels w/ intensity 1 = left-right; intensity 2 = front-back; intensity 3 = up-down

dir_image=$1
mask_image=$2

# create x,y,z images
${FSLDIR}/bin/fslmaths $mask_image -mul 0 arg0
${FSLDIR}/bin/fslmaths arg0 -add 1 arg

${FSLDIR}/bin/fslmerge -t xim arg arg0 arg0
${FSLDIR}/bin/fslmerge -t yim arg0 arg arg0
${FSLDIR}/bin/fslmerge -t zim arg0 arg0 arg

# get distance to each principal direction
echo “measuring…”
${FSLDIR}/bin/fslmaths ${dir_image} -abs tmpa
# x-dir
${FSLDIR}/bin/fslmaths tmpa -sub xim -sqr tmpx
${FSLDIR}/bin/fslmaths tmpx -Tmean -mul 3 tmpx
${FSLDIR}/bin/fslmaths tmpx -sqrt tmpx mv tmpx.nii.gz closex.nii.gz
# y-dir
${FSLDIR}/bin/fslmaths tmpa -sub yim -sqr tmpy
${FSLDIR}/bin/fslmaths tmpy -Tmean -mul 3 tmpy
${FSLDIR}/bin/fslmaths tmpy -sqrt tmpy mv tmpy.nii.gz closey.nii.gz
# z-dir
${FSLDIR}/bin/fslmaths tmpa -sub zim -sqr tmpz
${FSLDIR}/bin/fslmaths tmpz -Tmean -mul 3 tmpz
${FSLDIR}/bin/fslmaths tmpz -sqrt tmpz
mv tmpz.nii.gz closez.nii.gz

# combine and get index of closest direction echo
“making…”
${FSLDIR}/bin/fslmerge -t xyz closex closey closez
${FSLDIR}/bin/fslmaths xyz -mul -1 -Tmaxn -add $mask_image xyz

rm -rf closex.nii.gz closey.nii.gz closez.nii.gz tmpa.nii.gz arg.nii.gz arg0.nii.gz xim.nii.gz yim.nii.gz zim.nii.gz
echo “done!”
~~~

**Supplementary Figure S1.**
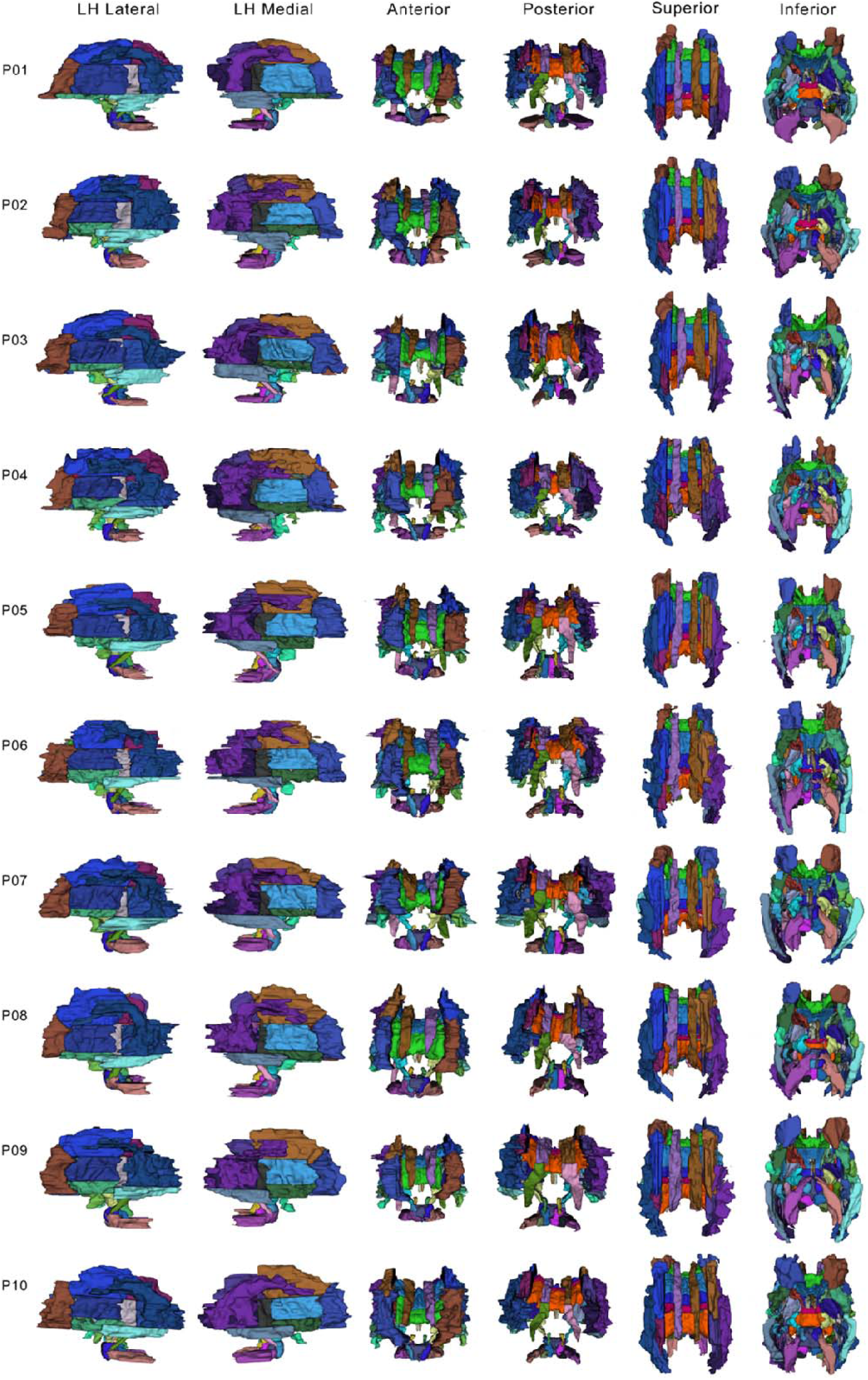
Surface representation of white matter (WM) parcellations for all 10 M-CRIB-WM participants.

**Supplementary Figure S2.**
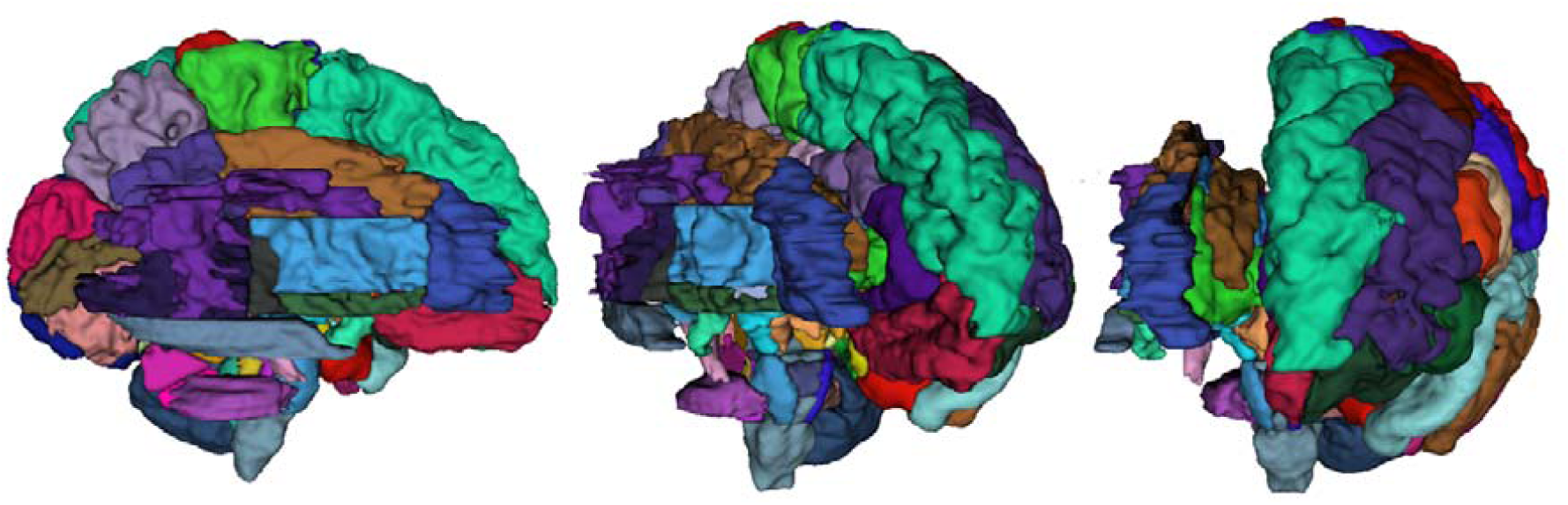
Combined surface representation of both M-CRIB 2.0 cortical parcellation (left hemisphere regions) and M-CRIB-WM white matter (WM) parcellations for a single participant.

